# Unveiling miR-451a and miR-142-3p as Prognostic Markers in NSCLC via sEV Liquid Biopsy

**DOI:** 10.1101/2024.11.29.622968

**Authors:** Miranda Burdiel, Ana Arauzo, Julia Jiménez, Rocío Moreno-Velasco, Carlos Rodríguez- Antolín, Olga Pernía, Oliver Higuera, Laura Gutiérrez-Sainz, Paloma Yubero, Julia Villamayor Sanchez, Itsaso Losantos-García, Nadina Erill Sagalés, Víctor González Rumayor, Javier de Castro, Inmaculada Ibáñez de Cáceres, Olga Vera

## Abstract

Despite advancements in personalized cancer therapies, platinum-based chemotherapy remains the cornerstone for treating solid tumors, including Non-Small Cell Lung Cancer (NSCLC). The integration of novel immunotherapies with platinum compounds has shown promising outcomes for the treatment of advanced disease. However, a significant proportion of patients experience therapeutic failure due to innate or acquired resistance. Thus, identifying molecular profiles and biomarkers to monitor patient progress and treatment response is crucial for tailoring therapeutic strategies. Small extracellular vesicle (sEV)-based liquid biopsy emerges as a promising non-invasive method for cancer management. sEVs play a critical role in cell communication and provide molecular insights into the tumor environment. In this study, we characterized the microRNome content of sEVs from cisplatin-resistant and -sensitive cancer cells using small-RNA sequencing. We identified and validated three miRNAs in two cohorts of 78 and 49 patients treated with either chemotherapy alone or chemo-immunotherapy, respectively, analyzed via liquid biopsy, differentiating NSCLC patients based on progression and overall survival. Notably, miR-451a emerged as a prognostic marker for chemo and chemo-immunotherapy, while miR-142-3p was identified for the first time as a potential prognostic marker specifically for stage IV patients, irrespective of the treatment. The combination of miR-451, miR-142-3p, and miR-55745, a novel miRNA identified from our miRNome screening, serves as a valuable biomarker for both cisplatin and chemo-immunotherapy treatment responses. This study underscores the role of sEVs in acquired cisplatin resistance and introduces novel miRNA-sEV biomarkers for managing NSCLC progression.

## INTRODUCTION

Non-small cell lung carcinoma (NSCLC) is one of the most frequent type of cancer worldwide (11,6% of new cases in 2018) and, by far, the most lethal (18,2% of all cancer-related deaths)^1^. The advanced stage of the disease at the time of diagnosis and the innate or acquired anti-cancer drug resistance are the main causes of this high mortality. Several mutations have been identified in lung cancer, such as in *EGFR*, *ALK*, ROS1, *BRAF, KRAS, MET exon 14 skipping, RET, NTRK* or *HER2*, that represent robust predictive biomarkers as well as very valuable therapeutic targets^2^. However, only between 15 to 30% of the cases have these targetable driven mutations^3^. Thus, in advanced stages with disseminated disease, which are not eligible for surgical resection, the standard treatment has been platinum-based chemotherapy. Currently, the new care-regime for advanced disease combines the use of platinum-based chemotherapy with novel immunotherapies. Unfortunately, there is a high percentage of patients who relapse from this treatment or do not respond initially, thus decreasing their overall survival. For this reason, it is crucial to identify new biomarkers for survival, prognosis, and drug-resistance in NSCLC, to be able to adapt the treatments to each patient. Exosomes are small-sized vesicles (30-150 nm) with an endosomal origin that are released by fusion of the multivesicular bodies with the plasmatic membrane^4^. These extracellular vesicles (sEVs) are released by most cell types as a mechanism of cellular communication, so they are found in different biological fluids such as blood, urine, saliva or cerebrospinal fluid^5–8^. sEVs are considered as “information shuttles” because they contain biologically active molecules such as proteins, RNA and DNA, which are capable of generating changes in the target cell after the internalization^4^. In fact, in the last decades numerous studies have shown the involvement of sEVs in diverse contexts, both physiological and pathogenic^8–10^. However, its role in cancer has gained special relevance in recent years, since tumor cells release a large amount of sEVs compared with normal cells^11^. Several studies describe their participation in tumorigenic processes such as immune system evasion, angiogenesis, premetastatic niche development or resistance to anti-cancer drugs^12–16^. Moreover, since the first report describing the presence of miRNAs in sEVs^17^, many researchers have focused their studies on sEVs-miRNAs as they can generate changes in a post-translational level. However, there is still a lack of prognostic markers identified and validated for their use in clinic. In the present study we show that high levels of sEV-miR-451a and miR-142-3p increase the risk of recurrence and death in NSCLC patients treated with platinum-based therapy alone or in combination with immunotherapy. Additionally, we have identified a novel miRNA, miR-55745, with potential as a biomarker in early in NSCLC. Our study provides useful biomarkers for the management of the advanced disease.

## MATERIAL AND METHODS

### Cell culture

Lung and ovarian cancer cell lines H23 and A2780 were purchased from the ATCC (Manassas, VA) or ECACC (Sigma-Aldrich, Spain). All of them were maintained in RPMI supplemented with 10% exosome-depleted FBS. FBS was depleted of bovine exosomes by ultracentrifugation at 100,000 × *g* for 16 h at 4°C. The CDDP-resistant variants H23R and A2780R were previously established by exposing cells to increasing doses of each platinum-based drug^18–21^. The CDDP sensitive and resistant ovarian cancer cell lines 41M and 41MR were provided by Dr. Kelland (UK) and maintained in DMEM supplemented with 10% exosome-depleted FBS.

### Clinical samples and data collection

Plasma samples from 78 NSCLC patients diagnosed with locally advanced and advanced stages (stages IIIA to IV) from 2015 to 2021 in La Paz University Hospital were collected before they received any platinum-based treatment. Additionally, we collected 49 plasma samples from NSCLC patients diagnosed with locally advanced and advanced stages (stages IIIA to IV) from 2019 to 2022 before they received platinum-based chemotherapy plus immunotherapy treatment. Follow-up was conducted according to the criteria of the medical oncology division of La Paz University Hospital. We also collected plasma samples from 18 healthy donors that comprised the control cohort. All the samples were processed following the standard operating procedures with the appropriate approval of the Human Research Ethics Committees, including informed consent within the context of research. All samples were processed within the first 30 minutes of collection using Vacutainer EDTA blood collection tubes in the case of blood and hemolyzed samples were discarded for the study as recommended^22^. Clinical, pathological and therapeutic data were recorded by an independent observer and blinded for statistical analysis.

### Exosome isolation

Cell line-derived exosomes, used for functional viability assays, were isolated from 500 ml of supernatant collected after 48-72 h of cell plating. Supernatant debris was pelleted by centrifugation at 1500 x *g* for 30 min. The supernatant was concentrated until obtaining 50 ml of each cell line using Ultra-15 Centrifugal Filter Concentrator (Merck, Germany) and filtrated with 0.2 µm filter to eliminate larger vesicles. Exosomes were then harvested by centrifugation at 100,000 × *g* for 2h. The exosome pellet was resuspended in 35 ml of 0.2 μm-filtered 1x PBS and collected by ultracentrifugation at 100,000 × *g* for 2 h (45ti rotor, Optima L-100 XP Centrifuge, Beckman Coulter, USA). Exosome pellets were resuspended in 200-300 µL of 0.2 μm-filtered 1x PBS and stored at -80°C. Cell line-derived exosomes, used for small RNAseq and proteomic analysis, were isolated by miRCURY Exosome Isolation Kit (Exiqon, Denmark) according to manufacturer’s instructions. Circulating exosomes and their miRNA content from plasma samples were obtained with exoRNeasy Serum/Plasma Midi Kit (Qiagen, Germany).

### Nanoparticle tracking analysis

Size and concentration of isolated sEVs samples from H23S, A2780S and 41M were characterized by Nanoparticle tracking analysis (NTA) using the NanoSight LM10 microscope equipped with NTA software v3.0 (Malvern, UK). Samples were diluted from 1:400 to 1:100 in 0.2 μm-filtered 1x PBS, depending on the sample concentration. Background extraction was applied and the automatic setting for minimum expected particle size, minimum track length and blur settings were employed. Three 30 s recordings at 30 frames per second were taken for each sample. The Nanosight automatic analysis settings were used to process the data.

### Exosome labelling and uptake assays by flow cytometry

Ultracentrifuge-obtained exosomes from culture medium of resistant cell lines were fluorescently labeled using PKH26 Red Fluorescent Cell Linker Mini Kit (Merk, Germany) according to the manufacturer’s protocol. Briefly, 250 µl of Dilutent C mixed with 1 µl of PKH26 were prepared for each sample. Exosome pellets were mixed with the stain solution and incubated for four minutes. The labelling reaction was stopped by adding an equal volume of 3% BSA 0.2 μm-filtered 1x PBS. Labelled exosomes were washed in 35 ml of 3% BSA 0.2 μm-filtered 1x PBS, collected by ultracentrifugation at 100,000 × *g* for 2 h and resuspended in PBS. Exactly the same process was performed with PBS as control to determine our PKH26 background.

For the uptake assays, sensitive cells were labelled with CellTrace Violet (CTV) (Thermofisher Scientific, USA). Briefly, 10^6^ cells resuspended in PBS + 5% exosome-depleted FBS were incubated 20 min with 20 µL of 1:100 CTV dilution. After washing the excess of dye with culture medium, cells were co-seeded with exosomes or PBS labelled with PKH26. After 20h of co-incubation, cells were trypsinized, washed and passed through the flow cytometer Facs Canto II (BD Biosciences, USA). To measure the death rate associated with the incubation with exosomes, we stained cells with 7-Aminoactinomycin D (7AAD) (BD Biosciences, USA). Results were analyzed using FlowJo (FLOWJO LLC, USA).

### sEVs functional assays

H23, A2780 and 41M sensitive cells were plated in 96-well plates in a concentration of 20,000 cells per well. sEVs isolated from resistant cells were quantified by Bradford assay (Bio-rad, CA, USA) and co-incubated with sensitive cells in a concentration of 80 µg/ml of sEVs for 48h followed by treatment with their respective Resistant-IC_50_ dose of cisplatin (Farma-Ferrer, Spain): 3.0 µg/ml for H23, 2.5 µg/ml for A2780 and 1 µg/ml for las 41M. Cell viability was measured 72 h after the drug treatment using CellTiter 96® AQueous One Solution Cell Proliferation Assay (MTS) (Promega, Spain). PBS-treated cells were used as control. Similarly, H23, A2780 and 41M resistant cells were plated in 24-well plates at a density of 40,000 cells per well and co-incubated with 80 µg/ml of sEVs or PBS for 48 h, followed by cisplatin treatment with their respective Sensitive-IC50: 0.5 µg/ml for H23, 0.25 µg/ml for A2780 and 1.5 µg/ml for 41M. After 72 h of treatment, cells were fixed with 1% glutaraldehyde (Merck, Germany) and stained with 0.1% crystal violet) (Merck, Germany). Dye was extracted with 10% acetic acid and absorbance was measured at 595nm using Infinite 200 PRO multimode reader (TECAN, Switzerland).

### Small RNAseq

Cisplatin-sensitive/resistant paired H23S/H23R, A2780S/A2780R and 41S/41R cells were used for the Small-RNAseq. RNA was extracted using a miRCURY RNA Isolation Kit -Cell and Plant (Exiqon, Denmark) according to manufacturer instructions. RNA was quantified with a NanoDrop ND-1000 spectrophotometer (Thermo Fisher Scientific, USA) and with a Qubit 4 fluorometer (Invitrogen, Thermo Fisher Scientific, USA), then analyzed with Arraystar (Arraystar Inc., USA) as follows: Total RNA of each sample was used to prepare the miRNA sequencing library; 3’ and 5’ adapter ligation, cDNA synthesis, PCR amplification and size selection between 130 and 150 bp (corresponding to 15[35 nucleotides (nt) of miRNAs) were performed. The DNA was sequenced with 51 cycles using the Illumina HiSeq 2000 sequencer (Illumina Inc., USA). The trimmed reads (length >= 15 nt and adapter removal) were aligned to human pre-miRNA in miRBase 21^1^ using NovoAlign software^2^ and miRNA read counts were normalized as tag counts per million (TPM) alignments^1^.

To select candidate miRNAs differentially represented between the sensitive and resistant phenotypes, reads with counts less than 2 were discarded. Next, they were normalized as transcripts per million aligned reads (TPM) and, to avoid false negatives, the miRNAs showing reads in both the sensitive and resistant subtypes were selected. For comparisons between phenotypes, the Log2 of Fold Change (log2FC) and p-value between each group were calculated. Next, those pre-miRNAs with a log2FC greater than or equal to 2 were selected. Finally, miRNAs meeting these conditions in at least 2 of the 3 lines analyzed were selected.

Novel or new microRNAs are those that are not currently annotated in the miRBase database but can be identified using prediction algorithms based on the counts obtained from massive miRNA sequencing. For this, miRDeep2^23^ was used on the final reads. For the selection of novel miRNAs, the same steps were followed as for known miRNAs, except for the last screening, in which those with a log2FC greater than or equal to 6 in any of the lines were selected.

### qRT-PCR

The nonspecific retrotranscription of all miRNAs from each sample was conducted using the TaqMan TM Advanced miRNA cDNA Synthesis Kit (Thermo Fisher Scientific, USA), according to manufacturer instructions. Quantitative analysis of each specific miRNA was performed using TaqMan Advanced miRNA assays (hsa-miR-151a-3p: 477919_mir; hsa-miR-451a: 478107_mir). A custom probe was designed by Thermo Fisher Scientific for hsa-miR-55745. TaqMan Universal PCR Master Mix (Thermo Fisher Scientific, USA) was used for qPCR amplification. All samples were analyzed in triplicate with a HT7900 Real-Time PCR System thermocycler (Applied Biosystems, USA) according to these settings: 10 min at 95°C and 40 cycles of 15 s at 95°C followed by 1 min at 60°C. The analysis of the results was performed with RQ Manager software (Thermo Fisher Scientific, USA) and relative miRNA levels were calculated according to the comparative threshold cycle method 2^-ΔCt^, where ΔCt is calculated by subtracting the Ct value of the endogenous control miR-151a^24^ from the Ct value of the targeted miRNA.

### Cell transfection and miRNAs functional assays

Each cell line was plated in 24-well plates at a density of 40,000 cells per well and incubated for 24h. Next, cells were transfected with 20nM of miR-451a (Ref: MC10286), miR-142-3p (Ref: MC10398) or negative control (Ref: 4464058) (Thermofisher Scientific, USA), using JetPrime (PolyPlus, France) and following the manufacturer’s protocol. Mimic for miR-55745 was synthesized based on its mature sequence by Thermofisher Scientific (Thermofisher Scientific, USA). After 6 h of transfection, cells were treated with increasing doses of cisplatin as previously reported^21^. After 72 h of treatment, cells were fixed with 1% glutaraldehyde (Merck, Germany) and stained with 0.1% crystal violet (Merck, Germany). Dye was extracted with 10% acetic acid and absorbance was measured at 595nm using Infinite 200 PRO multimode reader (TECAN, Switzerland). To validate miRNA overexpression, 200,000 cells plated in 6-well dishes were transfected with 20nM of each mimic for 72h followed by RNA isolation and quantification by qRT-PCR analysis.

### Statistical analysis

Patient’s clinical characteristics were described for the complete series with mean and standard deviation values or relative frequencies. The data were stratified for patients showing high or low levels of the miRNAs analyzed, and their distributions compared with the chi-squared test or Fisher’s exact test for qualitative variables, and Student’s t test or the Wilcoxon-Mann-Whitney test for quantitative variables. Overall survival and Progression free survival (PFS) were estimated according to the Kaplan-Meier method and compared between groups by means of the log-rank test. All the p-values were two-sided, and the type I error was set at 5 percent. A p-value below 0.05 was considered statistically significant. In vitro experiments were performed in triplicates or quadruplicates and each experiment was repeated at least twice. Unless otherwise indicated, one representative experiment is shown. Data represents the mean ± SD. All statistical analyzes were performed with SAS 9.3 (SAS Institute, Cary, NC, USA) and RStudio (version 1.1.423). Values of p<0.05 were considered as statistically significant, being * p<0.05, ** p<0.01 and *** p<0.001.

## RESULTS

### Exosomes from cisplatin-resistant cells confer drug resistance to sensitive cells

To study the involvement of sEVs in the process of cisplatin-resistance development, we co-cultured sEVs isolated from H23, A2780 and 41M CDDP-resistant cells (R-sEVs)^24^ with their CDDP-sensitive counterparts. First, we confirmed that R-sEVs were taken up by sensitive parental cells by labelling R-sEVs with PKH26, prior to exposing the sensitive cells to them. Using flow cytometry, we observed that uptake of sEVs by sensitive cells was successful in 61.2% for H23S (**Fig. 1A**), 75.6% for A2780S (**Fig. 1B**) and 46.3% for 41M (**Fig. 1C**) of alive cells when compared to PBS treated cells (**Fig. 1D**). Next, we exposed H23, A2780 and 41M sensitive (S), sensitive with R-sEVS (S+R-sEVs) and resistant (R) cells to the previously reported CDDP IC50 dose^3–5^. We observed that exposure of sensitive cells to R-sEVs increased their resistance to CDDP, being 1.7 times for H23, 2 times for A2780 and 1.6 times for 41M more resistant than the parental sensitive cell line, reaching the cell viability levels of the resistant cells in all cell lines tested (**Fig. 1E**). We also tested if sEVs from sensitive cells could revert the CDDP-resistant status. Thus, after confirming the isolation of sEVs from the media of H23S, A2780S and 41M cells (**Supplementary Fig. S1A**), we performed a co-culture experiment of S-sEVs with the resistant cells. We found that incubation of H23R, A2780R and 41MR with their respective sensitive sEVs did not alter the response to cisplatin (**Supplementary Fig. S1B**). Altogether, these results suggest that CDDP-resistant cells release biological molecules to the media through sVEs that can confer resistance to cisplatin.

**Figure 1.**
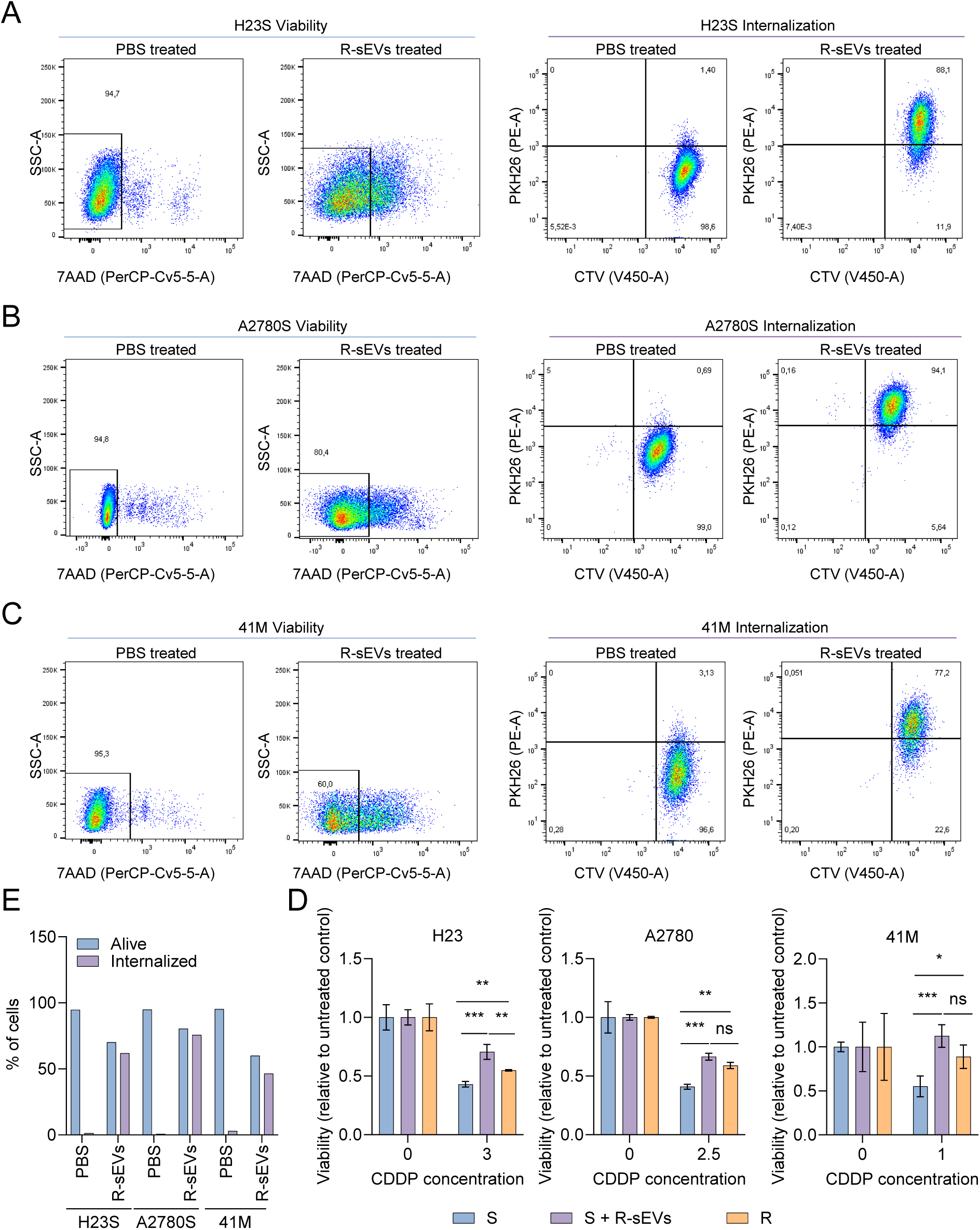
sEVs from CDDP resistant cells confer resistance to CDDP sensitive counterparts. (A-C) Flow cytometry assays to quantify the viability (left) and sEV internalization (right) performed in H23S (A), A2780S (B) and 41M (C) treated with either PBS or their resistant counterpart R-sEVs. (D) Quantification of sEVs internalization from (A-C). E) Cell viability assay of sensitive (S), sensitive with resistant-derived sEVs (S+R-sEVs) and resistant (R) cells after exposure to CDDP for 48h. One representative experiment out of two is shown. *, p<0.05; **, p<0.01; ***, p<0.001.

### sEV-miR-142-3p, miR-451a and miR-55745 arise as potential biomarkers of chemoresistance

Given that microRNAs are capable of inducing post-transcriptional changes, we focused on them to further investigate the molecular mechanisms underlying the acquisition of resistance in cisplatin-sensitive cells lines through sEVs derived from CDDP-resistant origin. Therefore, we performed a small RNA-seq on total RNA isolated from sEVs of the three matched CDDP-sensitive and resistant cell lines and compared their miRNA profile in order to identify candidates overrepresented in R-sEVs that could potentially induced resistance in sensitive cells. After the bioinformatics processing for the normalization of the results, we identified 603 and 506 miRNAs in H23S and H23R, 754 and 750 in A2780S and A2780R, and 725 and 790 in 41S and 41R, present in the sEVs content of the secretome (**Fig. 2A**). Next, we focused on common miRNAs showing up in both subtypes for each pair of cell lines, to discard those miRNAs in which there could be errors in the capture, obtaining 355 miRNAs for H23, 460 in A2780 and 499 in 41M (**Fig. 2A**). Then, we selected miRNAs overrepresented in resistant exosomes in comparison with the corresponding sensitive ones with a Fold Change > 2, identifying 16 miRNAs in H23, 2 in A2780 and 30 in 41M (**Fig. 2A** and **Supplementary Table S1**). To increase the success rate of our screening and find statistically robust candidates for subsequent clinical uses, we selected those miRNAs shared at least by two of the three lines, finally highlighting miR-892a, miR-891a-5p, miR-451a, miR-363-3p and miR-142-3p (**Fig. 2A**).

**Figure 2.**
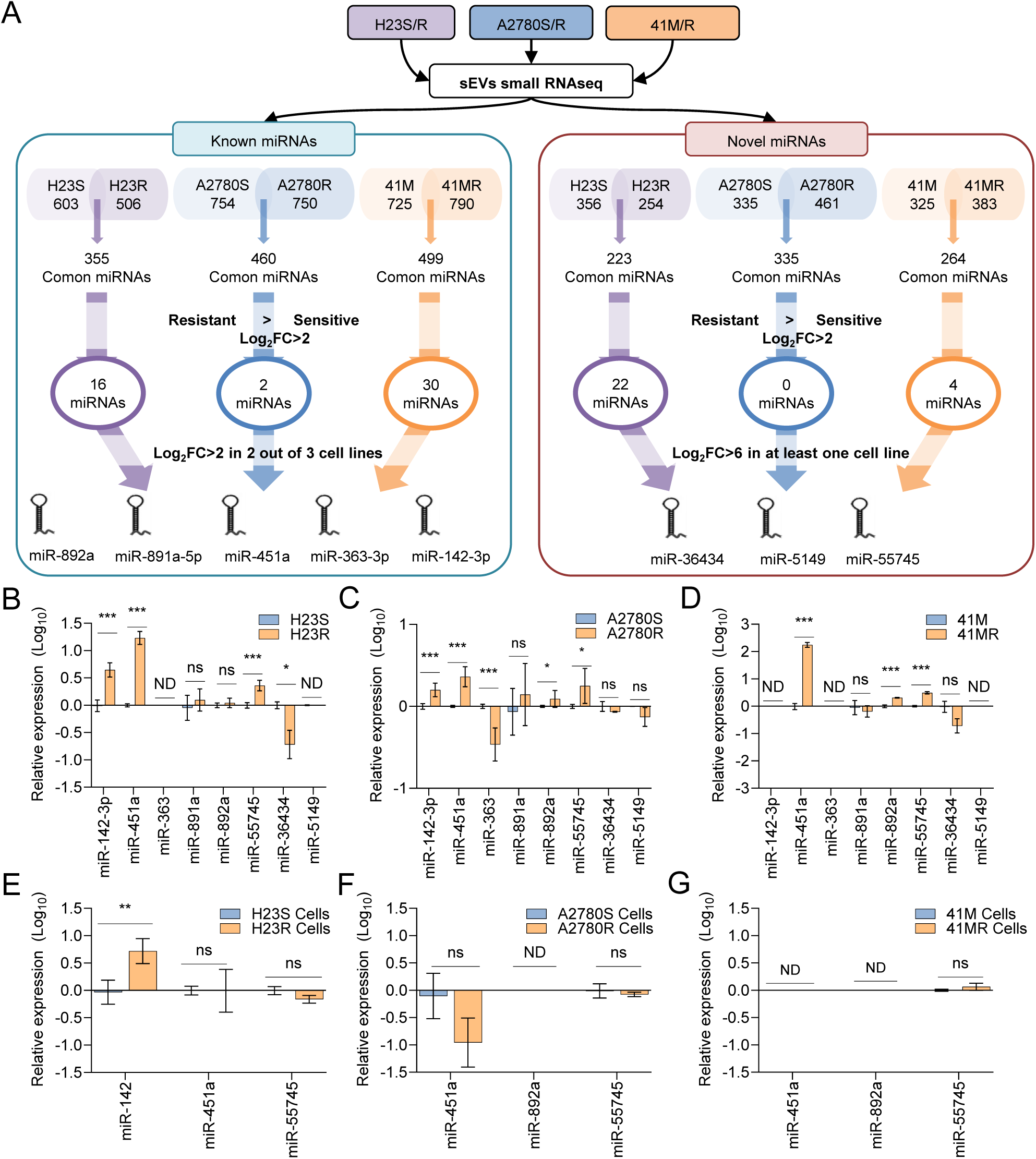
miR-142-3p, -451a and novel miR-55745 are potential sEVs biomarkers of chemoresistance in lung cancer. A) Pipeline for the identification of cisplatin-resistance sEV-miRNAs. Total RNA isolated from sEVs collected from cisplatin-Sensitive (S) and -Resistant (R) H23, A2780 and 41M, were subjected to small-RNA sequencing for the characterization of known (left) and novel (right) miRNAs as potential markers of cisplatin resistance. FDR and Log2FC were considered to select 8 miRNA candidates for further validation. (B) qRT-PCR analysis of the 8 candidate miRNAs in the sEVs from the cisplatin-Sensitive (S) and -Resistant (R) H23, A2780 and 41M. C) qRT-PCR analysis of miR-451a, -142 and -55745 as cisplatin-resistance candidate miRNAs in cisplatin-Sensitive (S) and -Resistant (R) H23, A2780 and 41M cells. One representative experiment out of two is shown.*, p<0.05; **, p<0.01; ***, p<0.001; ns, not significant; ND, not detected.

In parallel, the bioinformatics analysis of the small RNA-seq data allowed us to identify new miRNAs, not previously described in any current database (novel miRNAs). Following the same selection workflow, we identified 356 and 254 miRNAs in H23S and H23R, 335 and 461 in A2780S and A2780R, and 325 and 383 in 41S and R, respectively. Next, we obtained a total of 223 common miRNAs in H23, 335 in A2780 and 264 in 41, from which we selected those miRNAs with a Fold Change > 2 in resistant sEVs. This approach returned 22 miRNAs in H23, none in A2780 and 4 in 41 (**Fig. 2A** and **Supplementary Table S2**). Given that none of these miRNAs were shared by at least two of the cell lines, we added a more stringent filter and focused on those miRNAs with a Fold Change > 6 in at least one of the cell lines, identifying miR-novel-36434, miR-novel-5149 and miR-novel-55745 (names are assigned according with their chromosomal position) (**Fig. 2A**).

To validate the expression of these candidates with an alternative methodology, we performed qRT-PCR using Taqman probes. We evaluated changes in expression levels of the five known selected miRNAs and the three unknown miRNAs in the three matched cell lines using miR-151 as an endogenous control^24^. We found a significant increase of miR-451 and miR-55745 in sEVs of resistant vs sensitive cells from the three cell lines (**Fig 2B-2C**). Additionally, miR-142-3p was overrepresented in sEVs from H23R vs H23S (**Fig. 2B**) and A2780R vs A2780R (**Fig 2C**), being undetected in the cell line 41M (**Fig 2D**). We also found a significant increase of miR-892a in sEVs from resistant cells compared to sensitive ones for both ovarian cancer cell lines (**Fig. 2C, 2D**). To determine the potential relevance of these miRNAs intracellularly, we measured their basal expression levels in both the sensitive and resistant counterparts of H23, A2780 and 41M. We found that only miR-142-3p in H23R compared to H23S showed a significant expression increase (p<0.01) (**Fig. 2e**). Given that this was the only miRNA overexpressed in cells, and considering that both miR-451 and miR-55745 were validated in the three cell lines analyzed, we decided to move forward with these miRNAs to evaluate their potential as therapeutic-resistance biomarkers.

### Stage-specific sEV-miR-451a and sEV-miR-142-3p are biomarkers of chemotherapy-treated NSCLC patients

To determine the clinical relevance of our findings, we collected 78 plasma samples from advanced NSCLC patients (stages III and IV) who hadn’t received any treatment at the time of sample extraction. After that, all of them received platinum-based chemotherapy as first line of treatment and were clinically followed in order to monitor the time of relapse and overall survival. We isolated total sEVs-RNA and analyzed by qRT-PCR the levels of miR-451a, miR-142-3p and miR-55745. First, we compared the levels of these miRNAs in the NSCLC samples with the control ones (**Fig 3A-C**) and found that only miR-55745 levels were significantly lower in control plasma samples than in the lung cancer ones (**Fig. 3C**). This observation prompted us to consider whether higher or lower levels of miR-451a or miR-142-3p could differentiate groups of patients with worse prognosis. Thus, we divided patients according to the best cut-off for each miRNA, optimized through X tile^25^, into high levels and low levels (**Supplementary Table S3**) and compared with healthy donor samples. This approach showed that NSCLC with high levels of miR-451a (**Fig. 3D**), miR-142-3p (**Fig. 3E**), or miR-55745 (**Fig. 3F**), were significantly different than healthy donors, with no difference between low levels and controls (**Fig 3D-F**). Having observed that those patients with higher levels of miR-451a, miR-142-3p or miR-55745 were associated with NSCLC, we studied their implication in the pathology of the disease. We found a significant association between the levels of miR-451a and the histology type (p=0.012) and between miR-142-3p levels and NSCLC stage (p=0.012) (**Table 1**).

**Figure 3.**
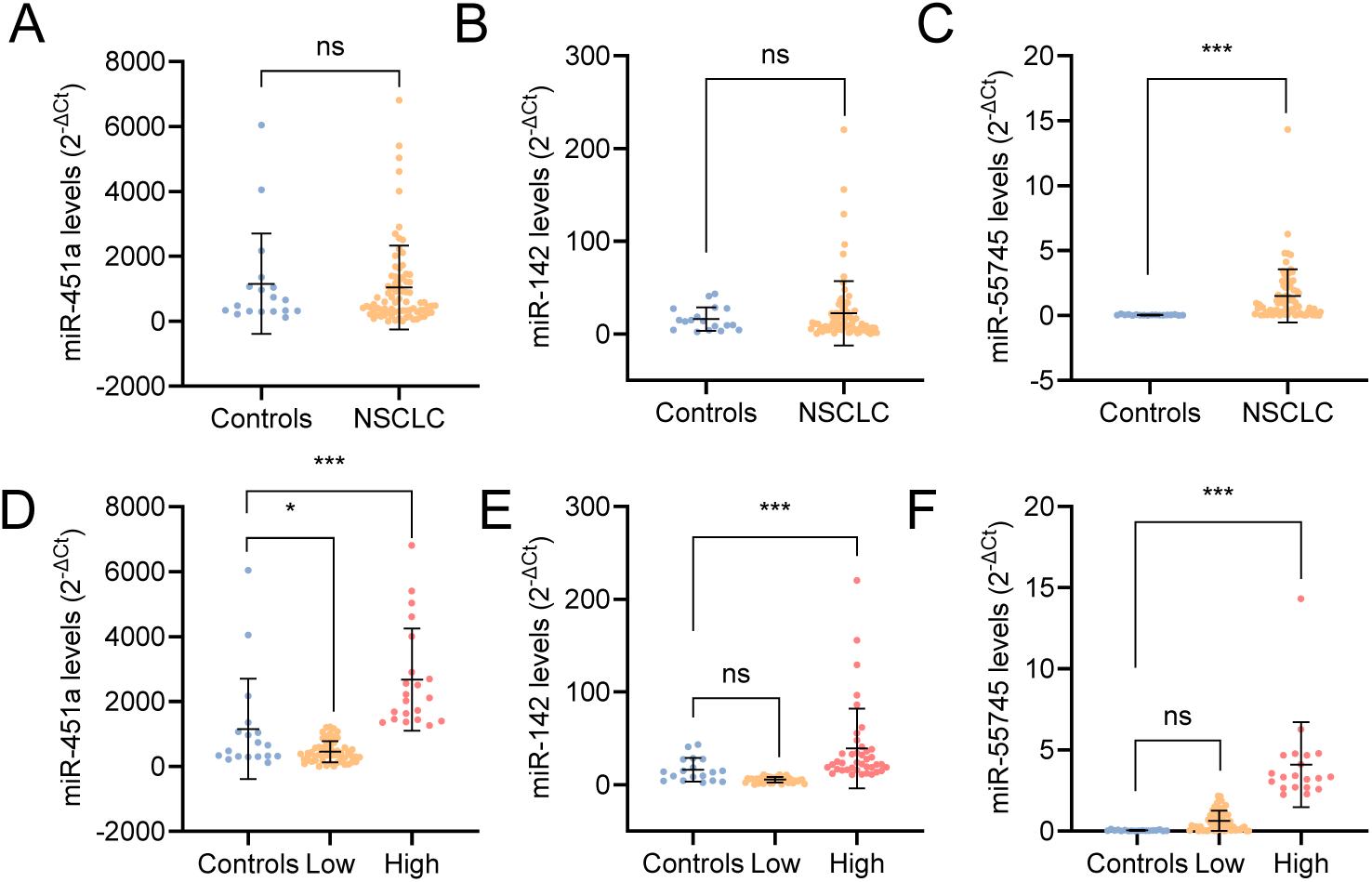
High levels of miR-142-3p, -451a and novel miR-55745 can differentiate lung cancer from control samples. (A-C) Levels of sEV-miR-451a (A), - miR-142-3p (B) and -miR-55745 (C) measured by qRT-PCR comparing healthy controls (n=18) and NSCLC patients (n=78). (D-F) Levels of sEV-miR-451a (D), -miR-142-3p (E) and -miR-55745 (F) measured by qRT-PCR comparing healthy controls (n=18) and NSCLC patients segregated into “low” and “high” levels according to the 75th percentile (miR-451a) or 50th percentile (miR-142-3p and miR-55745).*, p<0.05; ***, p<0.001; ns, not significant.

**Table 1.** Demographics of 78 Chemotherapy-treated NSCLC patients

|  |  | Complete<br>series<br>(n=78) | miR-451a<br>Low<br>(n=58) | miR-451a<br>High<br>(n=20) |  | miR-142-3p<br>Low<br>(n=40) | miR-142-3p High<br>(n=38) |  | miR-55745<br>Low<br>(n=59) | miR-55745<br>High<br>(n=19) |  |
| --- | --- | --- | --- | --- | --- | --- | --- | --- | --- | --- | --- |
|  |  | No. Of<br>patients | No. Of<br>patients | No. Of<br>patients | p | No. Of<br>patients | No. Of<br>patients | p | No. Of<br>patients | No. Of<br>patients | p |
| **Age (mean,**<br>**range)** |  | 67 (26-84) | 68 (47-84) | 64 (26-79) |  | 68 (48-84) | 65 (26-83) |  | 66 (26-84) | 67 (55-82) |  |
| **Sex** |  |  |  |  | 0.397 |  |  | 0.321 |  |  | 0.272 |
|  | Male | 56 | 39 | 17 |  | 28 | 28 |  | 38 | 18 |  |
|  | Female | 22 | 18 | 4 |  | 14 | 8 |  | 18 | 4 |  |
| **Smoker** |  |  |  |  | 0.575 |  |  | 0.903 |  |  | 0.108 |
|  | Active | 21 | 15 | 6 |  | 10 | 11 |  | 11 | 10 |  |
|  | Never | 4 | 4 | 0 |  | 2 | 2 |  | 3 | 1 |  |
|  | Inactive | 43 | 30 | 13 |  | 24 | 19 |  | 33 | 10 |  |
|  | NA | 10 | - | - |  | - | - |  | - | - |  |
| **Histology** |  |  |  |  | 0.014 |  |  | 0.802 |  |  | 0.657 |
|  | LUAD | 40 | 34 | 6 |  | 23 | 17 |  | 29 | 11 |  |
|  | LUSC | 28 | 15 | 13 |  | 14 | 14 |  | 21 | 7 |  |
|  | ND | 10 | 8 | 2 |  | 5 | 5 |  | 6 | 4 |  |
| **Stage** |  |  |  |  | 1 |  |  | 0.012 |  |  | 0.451 |
|  | III | 57 | 29 | 10 |  | 27 | 12 |  | 30 | 9 |  |
|  | IV | 21 | 28 | 11 |  | 15 | 24 |  | 26 | 13 |  |

Note: LUAD, Lung Adenocarcinoma; LUSC, Lung Squamous-cell carcinoma; NA, not available; ND, Not Defined.

Next, we evaluated the association between their expression levels and the progression free survival (PFS) and overall survival (OS) using Kaplan-Meier analysis. We found that high levels of sEVs-miR- 451a were significantly associated with lower PFS and OS (**Fig. 4A, 4B**), while patients with higher levels of miR-142-3p or miR-55745 were specifically associated with lower OS (**Fig. 4C, 4D**), with no differences for PFS (**Supplementary Fig. S2A, S2B**). Furthermore, high levels of miR-451a were associated with a 2.196-fold increased risk of progression (p=0.024; **Table 2**) and increased risk of death by 94.7% (p=0.026; **Table 2**). Similarly, elevated levels of miR-142-3p were linked to a 2.031-fold increased risk of death (p=0.012; **Table 2**). To gain clinically relevant insights, we analyzed the associations of these miRNAs to disease progression according to patient stages. Thus, using specific cut-off points for each stage (**Supplementary Table S3**) we observed that stage III patients with higher levels of miR-451a had a worse PFS (p=0.006; **Fig. 4F**) and OS (p=0.009; **Fig. 4E**), while no differences were found for the stage IV sub-cohort (**Supplementary Fig. S2C, 2D**). Conversely, higher miR-142-3p was specifically associated to worse OS only in stage IV patients (p=0.024; **Fig. 4G**), with no differences for stage III patients (**Supplementary Fig. S2E**). No differences were observed for miR-55745 in OS for either stage (**Supplementary Fig S2F, S2G**).

**Figure 4.**
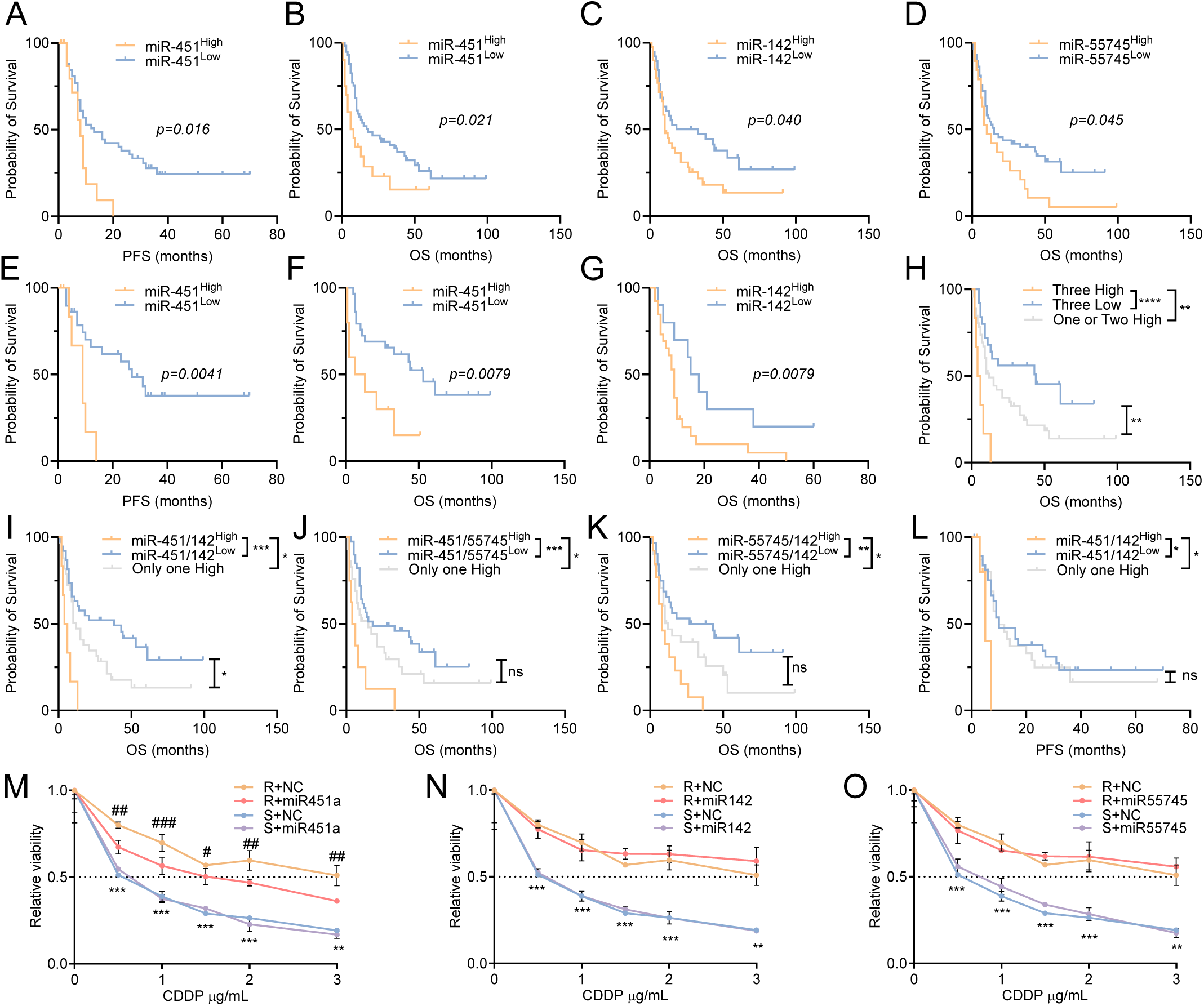
miR-451a is a prognostic sEVs biomarker of chemoresistance in advanced NSCLC cancer. (**A,B**) Kaplan-Meier survival analysis comparing Low (n=58) and High (n=20) miR-451a levels in NSCLC patients treated with chemotherapy in terms of progression free survival (A) and overall survival (B) in months. (**C**) Kaplan-Meier survival analysis of Low (n=40) and High (n=38) levels of miR-142-3p in terms of overall survival. (**D**) Kaplan-Meier survival analysis of Low (n=59) and High (n=19) levels of miR-55745 in terms of overall survival. (**E**) Kaplan-Meier survival analysis of Low (n=29) and High (n=10) miR-451a levels in stage III patients in terms of progression free survival. (**F**) Kaplan-Meier survival analysis of Low (n=29) and High (n=10) miR-451a levels in stage III patients in terms of overall survival. (**G**) Kaplan-Meier survival analysis of Low (n=11) and High (n=28) miR-142-3p levels in stage IV patients in terms of overall survival. (**H)** Kaplan-Meier survival analysis of Low (n=43), High (n=6) and “One or two high” (n=29) levels of a combination of miR-451a, -142 and -55745 in terms of overall survival. (**I**) Kaplan-Meier survival analysis of Low (n=34), High (n=13) and “Only one high” (n=31) levels of a combination of miR-451a and -142 in terms of overall survival. (**J**) Kaplan-Meier survival analysis of Low (n=44), High (n=9) and “Only one high” (n=25) levels of a combination of miR-451a and -55745 in terms of overall survival. (**K**) Kaplan-Meier survival analysis of Low (n=36), High (n=14) and “Only one high” (n=28) levels of a combination of miR-142-3p and -55745 in terms of overall survival. (**L**). Kaplan-Meier survival analysis of Low (n=34), High (n=13) and “Only one high” (n=31) levels of a combination of miR-451a and -142 in terms of progression free survival. Log-rank (Mantel-Cox) test was used for comparisons and p<0.05 was considered as a significant change in OS or PFS. *, p<0.05; **, p<0.01; ***, p<0.001; ****, p<0.0001. (**M-O**) Effect of overexpression of miR-451a (**M**), -142 (**N**) or - 55745 (**O**) on cell sensitivity to CDDP in H23 cell line. Viability curves of H23 transfected with negative control mimic (S+NC and R+NC) and with the overexpression mimics (S+miR-451a; R+miR-451a; S+miR-142-3p; R+miR-142-3p; S+miR-55745; R+miR-55745). Each experimental group was exposed for 48 h to six different CDDP concentrations, and data were normalized to each untreated control, set to 1. The data represent the mean ± SD of at least 2 independent experiments performed in triplicate at each drug concentration for each group analyzed. R vs S: **, p<0.01; ***, p<0.001; R+NC vs R+miRNA: #, p<0.05; ## p<0.01; ### p<0.001.

**Table 2.** Kaplan Meier and Cox regression analysis for miR-451a, miR-142-3p and miR-55745 in the chemotherapy-treated NSCLC cohort.

|  | PFS p-value | PFS HR (Low as Ref) | PFS IC 95% | PFS p-value HR | OS p-value | OS HR | OS IC 95% | OS p-value HR |
| --- | --- | --- | --- | --- | --- | --- | --- | --- |
| miR-451 | 0.016 | 2.196 | 1.109-4.348 | 0.024 | 0.021 | 1.947 | 1.083-3.499 | 0.026 |
| miR-142-3p | 0.756 | 1.430 | 0.804-2.542 | 0.224 | 0.04 | 2.031 | 1.170-3.529 | 0.012 |
| miR-55745 | 0.641 | 1.162 | 0.605-2.233 | 0.652 | 0.045 | 1.75 | 0.997-3.073 | 0.051 |
| Three high | 0.074 |  |  | 0.130 | 0.0001 |  |  | 0.001 |
| One or two High vs Three Low |  | 1.231 | 0.682-2.222 | 0.490 |  | 1.709 | 0.945-3.089 | 0.076 |
| Three High vs Three Low |  | 3.506 | 0.975-12.613 | 0.055 |  | 5.701 | 2.153-15.101 | 0.0001 |
| miR-451 & miR-142 | 0.112 |  |  | 0.125 | 0.0001 |  |  | 0.007 |
| One High vs Both Low |  | 1.592 | 0.858-2.951 | 0.140 |  | 2.299 | 1.250-4.230 | 0.007 |
| Both High vs Both Low |  | 2.382 | 0.957-5.850 | 0.058 |  | 2.773 | 1.270-6.058 | 0.011 |
| miR-451 & miR-55745 | 0.154 |  |  | 0.190 | 0.001 |  |  | 0.003 |
| One High vs Both Low |  | 1.428 | 0.787-2.590 | 0.241 |  | 1.656 | 0.939-2.920 | 0.082 |
| Both High vs Both Low |  | 2.48 | 0.840-7.322 | 0.1 |  | 4.015 | 1.772-9.093 | 0.001 |
| miR-142 & miR-55745 | 0.367 |  |  | 0.340 | 0.005 |  |  | 0.016 |
| One High vs Both Low |  | 1.070 | 0.570-2.010 | 0.832 |  | 1.377 | 0.741-2.558 | 0.311 |
| Both High vs Both Low |  | 1.739 | 0.834-4.082 | 0.131 |  | 5.864 | 1.456-5.864 | 0.003 |

Given the individual associations of these miRNAs with overall survival, we hypothesized that developing a miRNA scoring system might improve the prediction of mortality. Thus, we analyzed the combination of high vs. low levels of 1) all three miRNAs, 2) miR-451a and miR142, 3) miR-451a and miR-55745, and 4) miR-142-3p and miR-55745, based on their previous cut-offs and compared for survival analysis in the combined cohort, as this one showed global differences for the three miRNAs. We observed that the presence of high levels of these three miRNAs (**Fig. 4H**) were associated with lower OS and a 5.701-fold increased risk of death (p=0.0001; **Table 2**). Moreover, high levels of combined miR-451a and miR-142-3p levels (**Fig. 4I**), miR-451a and miR-55745 (**Fig. 4J**), or miR-142-3p and miR-55745 (**Fig. 4K**) were also associated with lower OS and increased risk of death in 2.773 (p=0.0001), 4.015 (p=0.001) and 5.864 times (p=0.003), respectively (**Table 2**). We also observed that high levels of combined miR-451a and miR-142-3p were significantly associated with lower PFS when compared to being both low (p=0.049) or only one high (p=0.047) (**Fig. 4L**). Finally, high levels of the three miRNAs also associated with a trend towards lower PFS (**Supplementary Fig. S2H**) compared to either three low (p=0.054) or to one or two high (p=0.043). No significant association with PFS or increased risk of relapse was found for the combination of miR-451a/miR-55745 or miR-142-3p/miR-55745 (**Supplementary Fig. S2I, 2J** and **Table 2**). To further characterize the role of these miRNAs in cisplatin response, we overexpressed miR-451a, miR-142-3p or miR-55745 in H23R and H23S cells using miRNA-mimics and tested their response to increasing doses of cisplatin. We found that miR-451 overexpression restored in part the cisplatin sensitivity in H23R (**Fig. 4M**), with no differences for miR-142-3p (**Fig. 4N**) or miR-55745 (**Fig. 4O**). Altogether these results indicate that miR-451a levels, and its combination with miR-142-3p are the best candidates for predicting worse clinical outcomes in chemotherapy-treated NSCLC patients.

### High sEV-miR-451a and –miR-142-3p alone and in combination are poor prognosis biomarkers in NSCLC patients undergoing chemo-immunotherapy

Given the current therapeutic approaches for advanced NSCLC^26^ and the promising results observed for these miRNAs regarding platinum-based therapy, we analyzed 49 plasma samples collected before starting chemo-immunotherapy treatment (CT-ICB). Clinical-pathological characteristics of this cohort (**Table 3**) were similar to the chemotherapy-treated one (**Table 2**) to the exception of the stage of the disease. However, similar to the chemotherapy-treated cohort, comparison of the levels of miR-451a, miR-142-3p, and miR-55745 with healthy donors (**Fig. 5A-5C**) showed that only levels of miR-55745 (**Fig. 5C**) were higher in the NSCLC patients than in the controls. To evaluate their potential as biomarkers, patients were stratified based on optimal cut-off levels for this cohort (**Supplementary Table S3**) and their levels were then compared to those in controls. We observed that plasma levels of miR-451a (**Fig. 5D**) and miR-142-3p (**Fig. 5E**) in the “low” group were significantly different than those in the healthy donors, with no differences in the “high” group. Moreover, levels of miR-55745 in the high group were significantly higher compared to the levels of this miRNA in the control samples (**Fig 5F**). Finally, our analysis revealed no significant association between the individual expression of these miRNAs and the clinicopathological data of the CT-ICB cohort (**Table 3**). This finding suggests that the prognostic potential of these miRNAs is independent of the initial clinical status of the patients. Kaplan-Meier survival analysis revealed that higher levels of miR-142-3p (**Fig. 5G**, **5H**) were significantly associated with worse PFS (**Fig. 5G**) and OS (**Fig. 5H**) in chemo-immunotherapy-treated patients, increasing the risk of relapse and death in 2.252 (p=0.047) and 2.696 (p=0.007) folds, respectively (**Table 4**). High levels of miR-451a were also associated with lower PFS (**Fig. 5I**) and OS (**Fig. 5J**) and increased the risk of relapse in 4.163 times (p=0.009) and death in 2.886 times (p=0.029) (**Table 3**). No association was found for miR-55745 (**Supplementary Fig. S3A, S3B**). Additionally, we evaluated the differences in OS and PFS between stage III and stage IV patients. For stage IV patients, we observed similar results (**Supplementary Fig. S3C-S3H**) to the combined analysis, likely due to the limited availability of stage III samples (n=8) and suggesting that the observed differences in the combined cohort were driven by stage IV patients. Therefore, we used the complete cohort to evaluate the predictive potential of combining miR-451a and miR-142-3p levels, since no differences were observed for miR-55745. We found that high levels of both miR-142-3p and miR-451a were linked to worse PFS (**Fig. 5K**) and OS (**Fig. 5L**) and increased the risk of relapse or death in 5.832 (p=0.004) and 5.512 times (p=0.004), respectively (**Table 4**). These results suggest that miR-142-3p and miR-451a levels can predict relapse and risk of exitus in the CT-ICB cohort.

**Figure 5.**
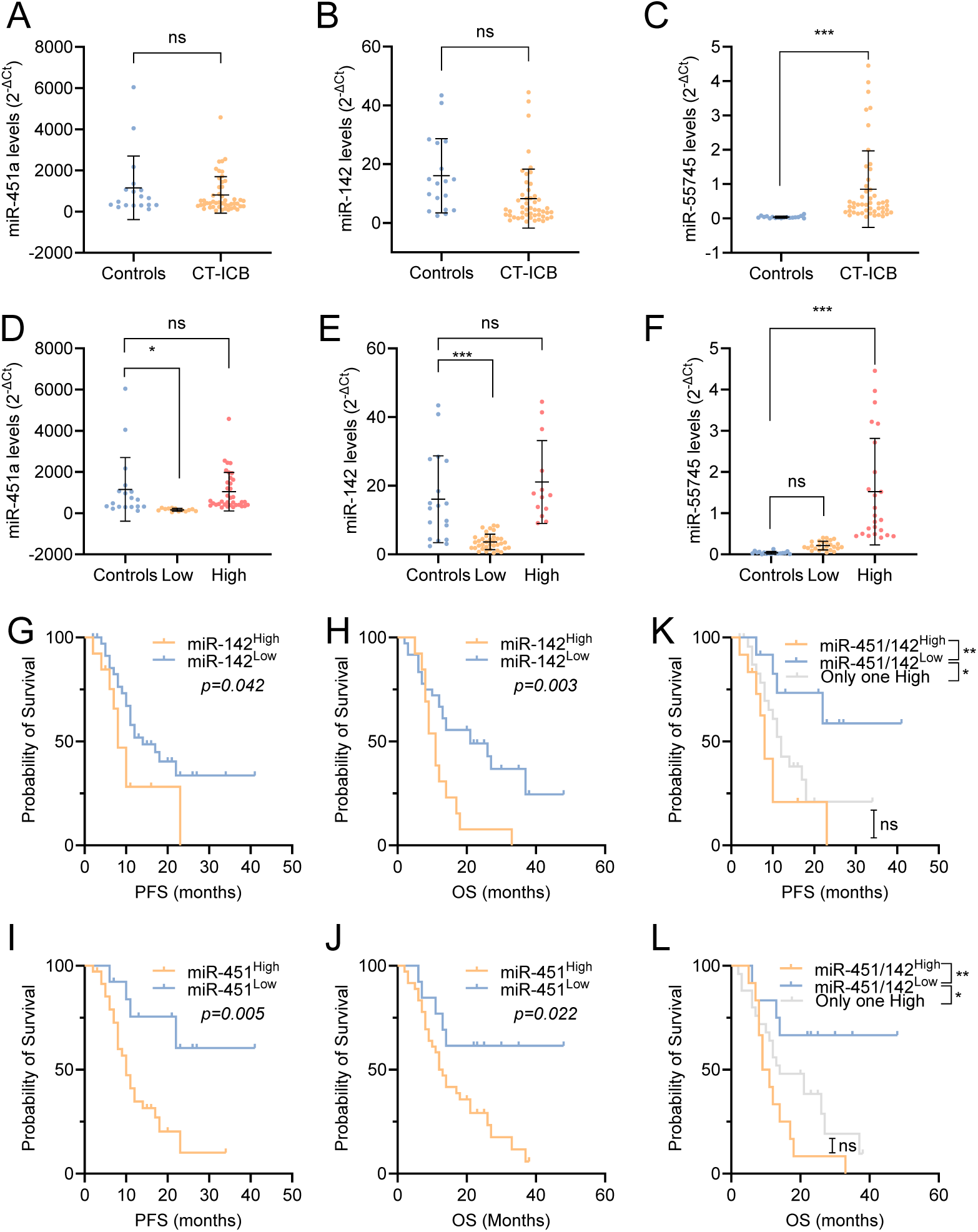
miR-142-3p and miR-451a are key predictors of chemo-immunotherapy resistance in advanced NSCLC. (**A-C**) Levels of sEV-miR-451a (A), -miR-142-3p (B) and -miR-55745 (C) measured by qRT-PCR comparing healthy controls (n=18) and NSCLC patients (n=49). (**D-F**) Levels of sEV-miR-451a (D), -miR-142-3p (E) and -miR-55745 (F) measured by qRT-PCR comparing healthy controls (n=18) and NSCLC patients segregated into “low” and “high” levels according to the 25^th^ percentile for miR-451a, 75^th^ percentile for miR-142-3p or 50^th^ percentile for miR-5574575.*, p<0.05; ***, p<0.001; ns, not significant. (**G,H**) Kaplan-Meier survival analysis of Low (n=36) and High (n=13) levels of miR-142-3p in terms of progression free survival (G) and overall survival (H). (**I,J**)Kaplan-Meier survival analysis of Low (n=14) and High (n=35) levels of miR-451a in terms of progression free survival (I) and overall survival (J). (**K,L**) Kaplan-Meier survival analysis of Low (n=12), High (n=16) and “Only one high” (n=12) levels of a combination of miR-451 and -142 in terms of progression free survival (K) and overall survival (L). Log-rank (Mantel-Cox) test was used for comparisons and p<0.05 was considered as a significant change in OS or PFS. *, p<0.05; **, p<0.01; ***, p<0.001; ****, p<0.0001.

**Table 3.** Demographics of 49 Chemo-immunotherapy-treated NSCLC patients

|  |  | Complete<br>series<br>(n=49) | miR-451a<br>Low<br>(n=13) | miR-451a<br>High<br>(n=36) |  | miR-142-3p Low<br>(n=36) | miR-142-3p High<br>(n=13) |  | miR-55745<br>Low<br>(n=25) | miR-55745<br>High<br>(n=24) |  |
| --- | --- | --- | --- | --- | --- | --- | --- | --- | --- | --- | --- |
|  |  | No. Of<br>patients | No. Of<br>patients | No. Of<br>patients | p | No. Of<br>patients | No. Of<br>patients | p | No. Of<br>patients | No. Of<br>patients | p |
| **Age (mean,**<br>**range)** |  | 65 (39-79) | 63 (43-79) | 65 (39-78) |  | 64 (39-79) | 65 (50-75) |  | 66 (43-75) | 64 (39-79) |  |
| **Sex** |  |  |  |  | 1 |  |  | 1 |  |  | 0.725 |
|  | Male | 39 | 10 | 29 |  | 29 | 10 |  | 19 | 20 |  |
|  | Female | 10 | 3 | 7 |  | 7 | 3 |  | 6 | 4 |  |
| **Smoker** |  |  |  |  | 0.326 |  |  | 0.669 |  |  | 0.348 |
|  | Active | 10 | 4 | 6 |  | 7 | 3 |  | 7 | 3 |  |
|  | Inactive | 31 | 6 | 25 |  | 24 | 7 |  | 15 | 16 |  |
|  | NA | 8 | 3 | 5 |  | 5 | 3 |  | 3 | 5 |  |
| **Histology** |  |  |  |  | 0.517 |  |  | 0.868 |  |  | 0.153 |
|  | LUAD | 29 | 2 | 20 |  | 22 | 7 |  | 18 | 11 |  |
|  | LUSC | 15 | 3 | 12 |  | 11 | 4 |  | 4 | 11 |  |
|  | ND | 5 | 1 | 4 |  | 3 | 2 |  | 3 | 2 |  |
| **Stage** |  |  |  |  | 0.523 |  |  | 0.389 |  |  | 0.367 |
|  | III | 8 | 3 | 5 |  | 7 | 1 |  | 4 | 4 |  |
|  | IV | 41 | 9 | 30 |  | 27 | 12 |  | 19 | 20 |  |

Note: LUAD, Lung Adenocarcinoma; LUSC, Lung Squamous-cell carcinoma; NA, not available; ND, not defined.

**Table 4.** Statistics associated to the levels of miR-451a, miR-142-3p and miR-55745 in the chemo-immunotherapy-treated NSCLC cohort.

|  | PFS p-value | PFS HR (Low as Ref) | PFS IC 95% | PFS p-value HR | OS p-value | OS HR | OS IC 95% | OS p-value HR |
| --- | --- | --- | --- | --- | --- | --- | --- | --- |
| miR-451 | 0.005 | 4.163 | 1.419-12.216 | 0.009 | 0.022 | 2.886 | 1.113-7.483 | 0.029 |
| miR-142-3p | 0.042 | 2.252 | 1.010-5.019 | 0.047 | 0.005 | 2.696 | 1.315-5.525 | 0.007 |
| miR-55745 | 0.838 | 0.926 | 0.441-1.943 | 0.838 | 0.388 | 1.344 | 0.685-2.640 | 0.39 |
| miR-451 & miR-142 | 0.008 |  |  | 0.015 | 0.007 |  |  | 0.012 |
| One High vs Both Low |  | 3.216 | 1.049-9.861 | 0.041 |  | 2.904 | 0.980-8.610 | 0.054 |
| Both High vs Both Low |  | 5.832 | 1.761-19.316 | 0.004 |  | 5.512 | 1.748-17.379 | 0.004 |

## DISCUSSION

In recent years, significant advances have been made in precision oncology, thanks to the molecular characterization of tumors and the study of their interactions with the tumor microenvironment using high-throughput analysis technologies. In this context, small extracellular vesicles represent a valuable source of tumor information, as they constitute a mechanism of cellular communication that is particularly enhanced in tumor cells^27^. In addition, these vesicles participate in different carcinogenic processes, allowing for a broader understanding of the molecular components responsible for the establishment and progression of tumorigenesis, thereby making them a promising source of biomarkers^27^.

In this study, we aimed to elucidate resistance mechanisms to platinum-based treatments in solid tumors and to identify novel biomarkers in liquid biopsies for non-small cell lung cancer (NSCLC) by examining miRNAs in circulating small extracellular vesicles (sEVs) from plasma in patients with this pathology. To achieve this, we conducted an initial experimental approach using a model based on three pairs of cisplatin-sensitive/resistant cell lines, allowing us to isolate and study the sEVs present in their secretome and to identify miRNAs with potential roles as CDDP-resistance biomarkers for subsequent translational validation in patient samples undergoing platinum-based combined treatments.

First, we optimized the experimental model for our study by determining the chemotherapy resistance induced by sEVs from the secretome of various cell models generated in our laboratory. Since 2012, numerous studies have demonstrated that sEVs from cells resistant to different antitumor drugs contain molecules capable of transmitting this resistance to neighboring cells, including breast, ovarian and lung cancers^28–34^. Our results demonstrate that sEVs derived from cisplatin-resistant cells can transfer part of their resistant properties to sensitive cells. Additionally, we observed that over 75% of the culture internalized sEVs after 20 hours of incubation, which is consistent with previous studies on sEV internalization efficiency in NSCLC^35,36^. Incubation of parental cells with sEVs from resistant cells resulted in basal mortality rates between 20% and 40%, depending on the cell line analyzed. This finding, which contrasts with other studies reporting minimal or no mortality in recipient cells^37^, may be attributed to the use of PKH26 as a marker, which could have cytotoxic effects compared to DiD used in other studies. Moreoer, higher sEVs doses might induce a stronger response in recipient cells, and variations in cell lines could also influence mortality rates^38^. We also evaluated the potential of sEVs from cisplatin-sensitive cells as a possible therapeutic target for drug sensitization. However, no change in the cisplatin response of resistant cells was observed. This could be due to the activation of drug resistance mechanisms independent of the active pathways in their respective parental cells after exposure to the aggressive chemotherapy agent. Studies using sEVs as therapeutic tools focus on their use as vehicles to deliver drugs or interfering RNAs to target cells, demonstrating the inactivation of new processes that had been established in the pathology^39^.

After selecting the paired cell lines H23, A2780, and 41M as a model to continue our study, we compared the sEVs miRNA levels between the sensitive and resistant phenotypes of each pair using massive miRNA sequencing analysis (small RNA-seq) to identify new candidates involved in platinum derivative resistance mechanisms. In recent years, the study of the sEVs microRNome has gained substantial prominence, reflecting the growing interest in understanding the role of microRNAs in small extracellular vesicles (sEVs). Our study adds to this expanding field by employing advanced sequencing techniques, having allowed us to identify around 300 novel miRNAs and 500-700 known miRNAs in each pair of cell lines, thereby contributing valuable insights to the current knowledge. Out of the identified miRNAs, we selected the eight most over-represented in the sEV content from resistant cells, being three of them non-described yet. Following the criteria described by miRBase, we used quantitative PCR as an alternative technique to validate the expression differences identified by NGS and obtained a coincidence rate between the two techniques of approximately 50%, a success rate consistent with the results obtained in a previous study conducted with NSCLC sEVs^40,41^. A total of four miRNAs were validated, miR-451a, miR-142-3p, miR-892a-5p and the novel miR-55745. For some miRNAs, the amplification obtained by qRT-PCR was very low or non-existent, as was the case for the novel miR-5149 in all cell lines or miR-363-3p and miR-142-3p in 41M/MR. This discrepancy might be attributed to the sEV levels of these candidates being low yet detectable through the sensitivity of massive sequencing techniques, while posing amplification challenges for PCR. Our findings highlight that while massive miRNA sequencing is a powerful tool for studying numerous miRNAs simultaneously, it is crucial to validate these results using well-established, robust techniques like qRT-PCR. To establish the role of these miRNAs as potential liquid biopsy biomarkers, we analyzed their levels in a cohort of 78 advanced-stage NSCLC patients treated with platinum-based compounds and a cohort of 49 advanced-stage NSCLC patients treated with a combination of platinum-based and immunotherapy. Both cohorts were well standardized by histological types, male/female ratio, treatments, and associated survival data. We observed an overrepresentation of stage III cases in the chemotherapy-treated cohort and stage IV cases in the chemo-immunotherapy-treated cohort, consistent with current management recommendations for these patients^42^. Given that samples were obtained before any treatment, this imbalance in stages could explain the differences observed in miRNA levels in NSCLC samples compared to control samples.

Among the three miRNAs analyzed, miR-55745 demonstrated the lowest predictive potential, aligning with our functional assays using mimics, where no changes in cell viability to cisplatin were observed. Segregating patients into “low” and “high” miR-55745 level groups revealed a slight difference in overall survival but had no effect on disease-free survival in the platinum-treated cohort.

Similarly, we observed non-significant results in the cohort treated with a combination of platinum and immunotherapy. These findings, supported by our in vitro experiments overexpressing miR-55745, suggest that miR-55745 may not play a role in cisplatin resistance. However, given the observed differences between NSCLC and control samples, a potential role in the early stages of NSCLC cannot be completely discarded and future studies will address this.

miR-451a, whose increased levels in resistant-sEV were validated by qRT-PCR in the three analyzed cell lines, is widely recognized in the literature as a tumor suppressor miRNA in various types of cancer^43^. In glioma cells, miR-451a inhibits growth through the regulation of the PI3K/AKT pathway via CAB39^44^. Additionally, the negative regulation of this miRNA has been linked to resistance to antitumor drugs in breast cancer, renal cancer and NSCLC^45–48^. Notably, one study in breast cancer observed increased sensitivity to doxorubicin after transfection of MCF-7 cells with miR-451a^49^, mirroring our findings where resistant phenotype cells showed increased sensitivity to cisplatin after miR-451a transfection. Therefore, the absence of miR-451a expression in the cellular content of any analyzed cells, coupled with its established role as a tumor suppressor, suggests that resistant cells may expel it via sEVs as a survival mechanism, a hypothesis previously postulated^50,51^. Our findings suggest that miR-451a plays a complex, context-dependent role in lung cancer, influencing survival differently across stages and treatments. In stage III patients treated solely with chemotherapy, elevated miR-451a levels significantly correlate with worse prognosis. This relationship may stem from miR-451a’s role in promoting local tumor progression by regulating oncogenic pathways such as KIF2A^52^. In contrast, in stage IV patients, where metastases are established, this effect may be diluted by the more heterogeneous tumor microenvironment and greater tumor burden, with other factors playing a more dominant role. Additionally, miR-451a might be critical during the transition to metastasis but less influential in maintaining metastatic states. Interestingly, in the cohort receiving chemotherapy combined with immunotherapy, high miR-451a levels correlate with poorer OS and PFS, even in stage IV patients, suggesting a potential immunomodulatory role. Indeed, miR-451a is involved in various pulmonary conditions—such as acute lung injury, bronchopulmonary dysplasia, tissue-engineered vascular graft stenosis, and asthma—through mechanisms like macrophage migration and activation^53^. This aligns with findings from other diseases, where miR-451a has been shown to influence immune responses by attenuating innate immune activity in macrophages and dendritic cells^54^ or by enhancing the phagocytic activity and differentiation of macrophages^55^. Moreover, miR-451a can affect T-cell populations and cytokine profiles in systemic lupus erythematosus, underscoring its broad immunoregulatory functions^56^. Although no studies have yet explored miR-451a’s role in lung cancer immunotherapy, its association with immune responses in other contexts is compelling. In melanoma, for example, elevated miR-451a levels have been linked to better responses to anti-PD-1 treatment^53^. These findings highlight the potential relevance of miR-451a in modulating the tumor immune microenvironment in lung cancer. Further research could uncover its role as a biomarker for predicting immunotherapy responses or as a therapeutic target to improve treatment outcomes.

The overrepresentation of miR-142-3p in sEVs from cisplatin-resistant lung cancer cells suggested that it could be a resistance mechanism to platinum derivatives in this tumor type. The role of this miRNA is controversial; numerous studies describe it as a tumor suppressor miRNA, as it is strongly inhibited in various types of cancer and negatively regulates known oncogenes^57–60^. It has also been associated to chemoresistance through the regulation of HMGB1 in acute myeloid leukemia and NSCLC^61,62^. Conversely, other studies point to its involvement as an oncogenic miRNA in colorectal cancer, renal carcinoma and gastric cancer^63,64^ through the regulation of the transcription factor FOXO4. As a sEV-miRNA, its dual nature as a tumor suppressor miRNA and oncomiR capable of promoting carcinogenesis in both the cell that expels it and in target cells has also been described^50^. Our functional assays with mimics showed no differences in cisplatin viability of sensitive or resistant cells transfected with miR-142-3p. This result aligns with our plasma-derived sEVs analysis, where high miR-142-3p levels specifically associate with stage IV NSCLC patients, which could help predict which patients are at risk of developing metastatic disease. Moreover, miR-142-3p levels show a strong association with poorer prognosis and increased relapse in platinum-immunotherapy treated patients. Importantly, recent studies show a role for miR-142-3p as a negative regulator of the anti-tumor immunological response^65–67^. Future studies involving co-culturing experiments will determine the actual role of miR-142-3p in reducing the response to immunotherapy in NSCLC.

Finally, our experimental approach reveals a potential synergy between miR-142-3p and miR-451a, as patients with elevated levels of both sEV-miRNAs exhibited a five-fold decrease in progression-free survival and overall survival across both cohorts. Given that both miR-451a and miR-142-3p negatively regulate important cell proliferation and survival pathways such as RAF/MEK/ERK^68,69^ and PI3K/AKT^62,70^, their cell secretion through sEVs, as recently shown for miR-451a^51^, could promote mechanisms associated with tumor proliferation and drug resistance in NSCLC. These findings suggest that alternative treatments, including specific inhibitors targeting these pathways, might benefit patients with elevated levels of both miRNAs in the sEV-content who do not respond to first-line platinum treatment.

Altogether, our study demonstrates that plasma-derived sEV levels of miR-451a and miR-142-3p, either individually or in combination, serve as valuable prognostic biomarkers and predictors of platinum response in liquid biopsy for advanced-stage NSCLC patients treated with chemotherapy or chemo-immunotherapy. These findings highlight the potential clinical utility of these miRNAs in enhancing personalized treatment strategies and improving patient outcomes.

## DECLARATIONS

### Ethics approval and consent to participate

All samples were processed following the standard operating procedures with the appropriate approval of the Human Research Ethics Committees, including informed consent within the context of research (HULP: PI-3508 and -5063).

### Availability of data and materials

The datasets generated and/or analyzed during the current study are available in the GEO repository, number GSE204944.

### Authors contributions

Conceptualization, MB, IIC. Methodology, MB, AA, JJ, RM, CRA, OP. Formal analysis: MB, AA, JJ, RM, CRA, OP, ILG, IIC, OVP. Investigation, MB, AA, JJ, RM, CRA, OP, OH, LGS, PY, ILG, NES, VGR, JDC, IIC, OVP. Resources OH, LGS, PY, ILG, NES, VGR, JDC, IIC, OVP.

Writing original draft, JJ, OV, IIC. Writing, review and editing, MB, AA, JJ, RM, CRA, OP, OH, LGS, PY, ILG, NES, VGR, JDC, IIC, OVP. Supervision, OV, IIC. Project administration, OVP, IIC. Funding acquisition, NES, VGR, JDC, IIC.

### Authorship

We declare that all the authors of this study have directly participated in the planning, execution, or analysis of the study, and all the authors have read and approved the final version submitted, adhering to the guidelines of the ICMJE.

## Acknowledgments

The authors thank HULP-IdiAPZ Biobank for sample processing, the cell culture, Flow Cytometry and Microscopy Cores at IdiPAZ. We also thank the financial support.

