## Supplementary Figures for "Unveiling miR-451a and miR-142-3p as Prognostic Markers in NSCLC via sEV Liquid Biopsy"

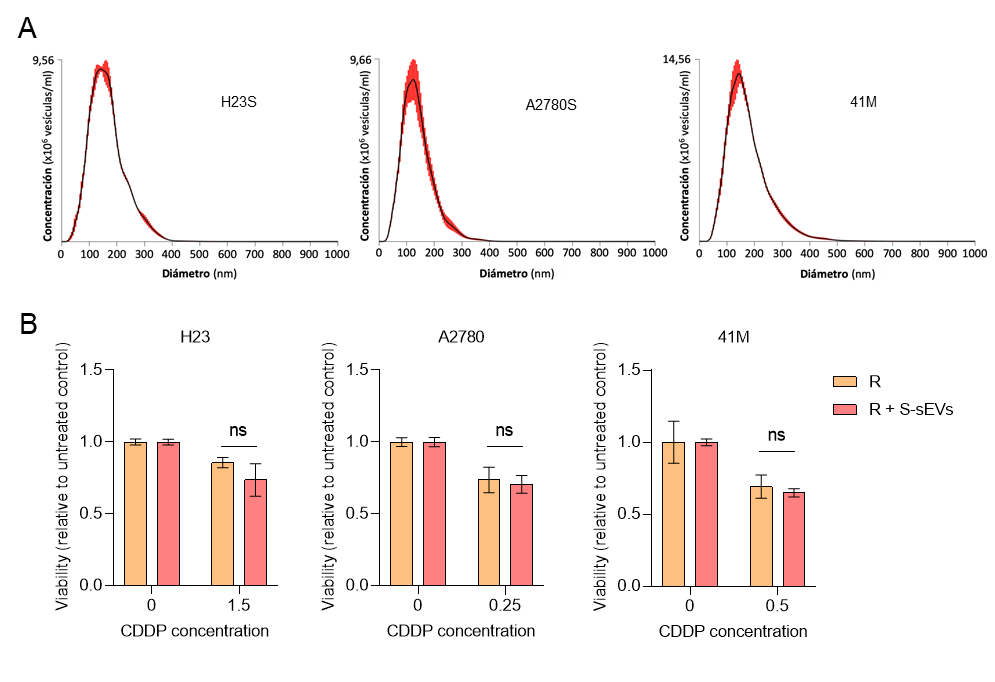

**Supplementary Figure 1.** A) Nanoparticle Tracking Analysis of sEVs isolated from H23R, A2780R and 41MR. B) Cell viability assay of Resistant (R) and resistant with sensitive-derived sEVs (R+S-sEVs) cells after exposure to CDDP for 48h. One representative experiment out of two is shown. Ns, not significant.

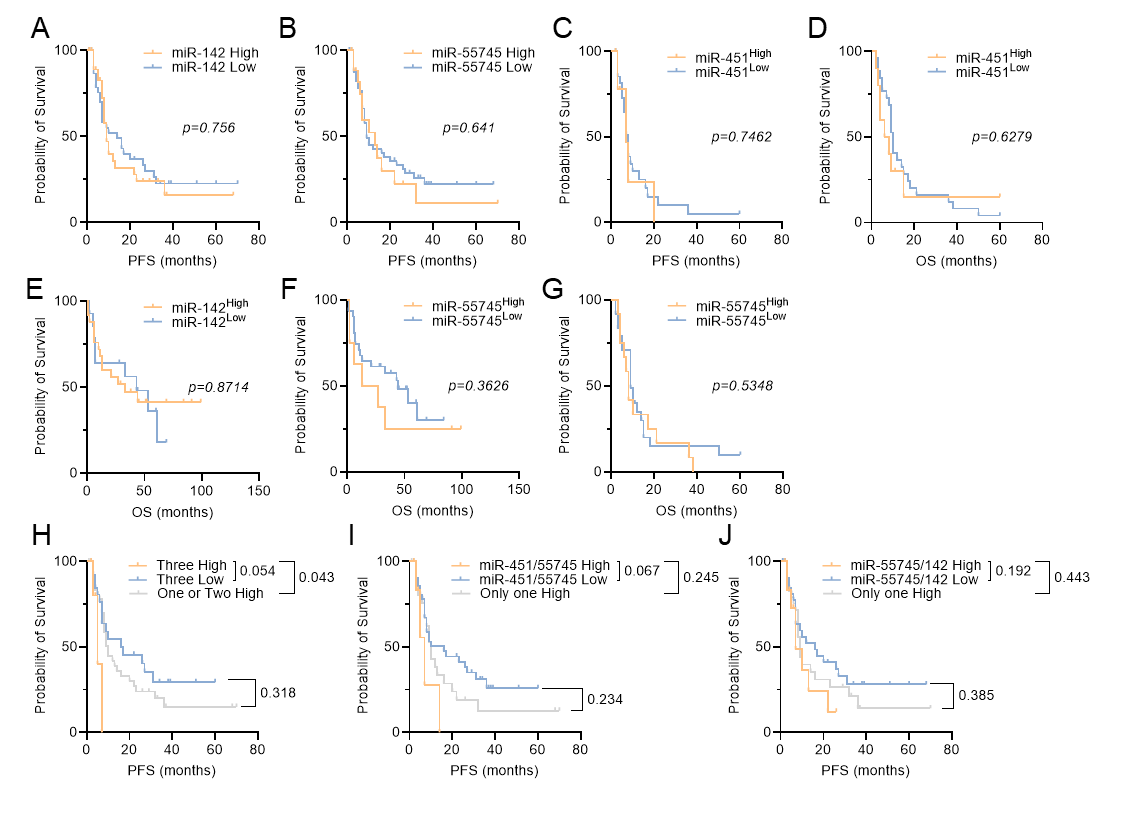

**Supplementary Figure 2.** (**A**) Kaplan-Meier survival analysis of Low (n = 40) and High (n=38) levels of miR-142-3p in terms of progression free survival. (**B**) Kaplan-Meier survival analysis of Low (n = 59) and High (n=19) levels of miR-55745 in terms of progression free. (**C,D**) Kaplan-Meier survival analysis of Low (n=29) and High (n=10) miR-451a levels in stage IV patients in terms of progression free survival (**C**) and overall survival (**D**). (**E**) Kaplan-Meier survival analysis of Low (n=15) and High (n=24) miR-142-3p levels in stage III patients in terms of overall survival. (**F,G**) Kaplan-Meier survival analysis in Stage III patients comparing Low (n=30) and High (n=9) miR-55745 levels (F) and in Stage IV patients comparing in terms of Low (n=26) and High (n=13) miR-55745 levels in terms of overall survival (G). (**H**) Kaplan-Meier survival analysis of Low (n=43), High (n=6) and “One or two high” (n=29) levels of a combination of miR-451a, -142 and -55745 in terms of progression free survival. (**I**) Kaplan-Meier survival analysis of Low (n=44), High (n=9) and “Only one high” (n=25) levels of a combination of miR-451a and -55745 in terms of progression free survival. (**J**) Kaplan-Meier survival analysis of Low (n=36), High (n=14) and “Only one high” (n=28) levels of a combination of miR-142-3p and -55745 in terms of progression free survival. Log-rank (Mantel-Cox) test was used for comparisons and p<0.05 was considered as a significant change in OS or PFS.

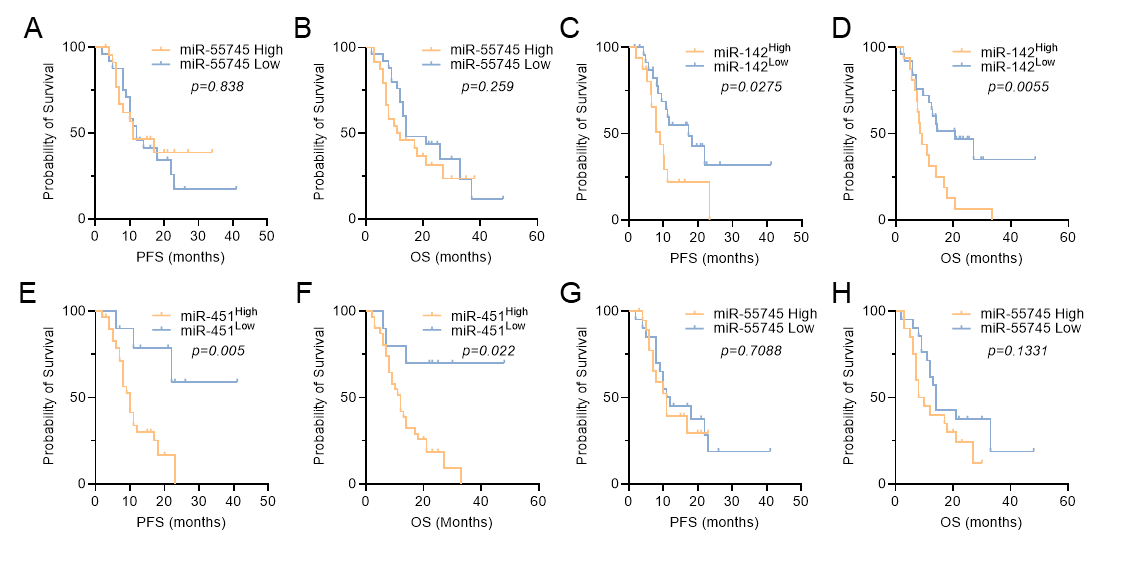

**Supplementary Figure 3.** (**A,B**) Kaplan-Meier survival analysis of Low (n = 24) and High (n=25) levels of miR-55745 in terms of progression free survival (A) and overall survival (B). (**C,D**) Kaplan-Meier survival analysis comparing Low (n=25) and High (n=16) miR-142-3p levels in stage IV NSCLC patients in terms of progression free survival (**C**) and overall survival (**D**) in months. (**E,F**) Kaplan-Meier survival analysis comparing Low (n=10) and High (n=31) miR-451a levels in stage IV NSCLC patients in terms of progression free survival (**E**) and overall survival (**F**) in months. (**G,H**) Kaplan-Meier survival analysis comparing Low (n=21) and High (n=20) miR-451a levels in stage IV NSCLC patients in terms of progression free survival (**G**) and overall survival (**H**) in months.
